# Intergenerational transmission of parental rejection effects on survival, behaviour, and miRNA expression in owl monkeys

**DOI:** 10.1101/2024.12.20.629709

**Authors:** Jayde Farinha, Mirtha Clausi, Fabian Orlando Chamba Pardo, Nofre Sanchez-Perea, James Boot, Chaz Mein, Ping Yip, Ursula M. Paredes

## Abstract

Parental rejection in captive primates represents severe early-life stress, yet intergenerational effects remain poorly characterised. We investigated spontaneous rejection in owl monkeys (*Aotus nancymaae*) across two generations using demographic records (n=1,372), behavioural observations (n=65), and blood miRNA profiling (n=47). Rejected individuals showed 51% reduced lifespan, decreased probability of reaching reproductive age, and increased morbidity throughout life. Critically, well-reared offspring of rejected parents exhibited similar reductions in survival to reproductive age and lifespan. Rejected adults displayed heightened fearfulness and aggression with reduced paternal care; offspring showed decreased locomotion and altered stress responses. Blood miRNA analysis revealed significant upregulation of mml-miR-30a-5p in female offspring-a miRNA associated with early-life stress and behavioural disorders in humans. These findings demonstrate intergenerational transmission of rejection effects on fitness and behaviour, with preliminary evidence suggesting epigenetic mechanisms are associated with this transmission, though multiple pathways likely operate.

## Introduction

Epidemiological research in humans increasingly suggests that stress experienced by parents can be transmitted to offspring, affecting their health and behaviour across the lifespan^1–4^. However, studying parental stress and its intergenerational effects in humans is challenging due to long lifespans and complex social confounds; non-human primates offer valuable model systems due to their shared cognition, physiology, and extended parental investment with humans.

Studies in primates exposed to early-life adversity (ELA), including experimental parental separation, harsh parenting, or parental loss, demonstrate that these individuals exhibit lifelong alterations in stress behaviour and neural circuits^5–7^, immune function^8^, and health outcomes^9^. Critically, ELA is associated with decreased survival in early life and reduced lifespan in the wild^10,11^ and in free-ranging populations^12^.

Complete parental rejection, e.g. when mothers refuse to nurse newborns, represents the most severe form of ELA, as it can be lethal without intervention^13^. Rejection is a persistent problem in captive primate colonies globally, documented across the primate order^14–26^. Veterinarians typically intervene with hand-rearing or nursery-rearing to prevent infant mortality, and whilst rejected-then-reared primates are known to behave abnormally and show physiological differences^5,13^, the long-term effects on health and life history remain poorly characterized outside a handful of species.

Emerging evidence suggests some effects of stressful rearing experience are heritable: offspring of ELA-exposed parents show altered behaviours and immune profiles^27–29^ in captive macaques and reduced longevity and survival^30^ in wild baboons. If rejection were to affect the health and behaviour of rejected and rescued captive primates, their descendants could also suffer. Studying intergenerational transmission of stress in captive populations can shed light on underlying mechanisms and ultimately inform strategies to improve captive primate welfare.

Candidate mechanisms for intergenerational transmission of stress, including maternal neglect, suggest alterations in epigenetic marks that become modified in response to environmental cues^31^. In captive primates, altered DNA methylation patterns in stress-regulatory genes are associated with ELA and predict health outcomes^32–34^.

More recently, circulating microRNAs (miRNAs) have emerged as potential carriers of intergenerational stress. In *C. elegans*, small RNAs can be inherited across multiple generations and mediate transgenerational responses to environmental stress^35^. Altered miRNA expression has been detected in stressed mammals and their offspring, including in sperm^36^ and blood plasma^37^. Multiple pathways may contribute to intergenerational transmission, including germline epigenetic inheritance, *in utero* programming through maternal physiology, or subtle alterations in parental behaviour that affect offspring development.

We hypothesized that if complete parental rejection is associated with behavioural effects detectable in the next generation, transmission might be accompanied by alterations in epigenetic marks such as miRNAs. We tested this in a captive population of owl monkeys (*Aotus nancymaae*)—a Neotropical primate that is uniquely monogamous, nocturnal, and displays intensive biparental care^38,39^. In this colony, spontaneous parental rejection occurs in approximately 5-10% of births. Mothers refuse nursing, occurring primarily within the first days or months after birth, and occasionally paternal aggression accompanies maternal rejection. Nursery-rearing is practiced to rescue rejected infants.

We previously reported poor health outcomes in rejected juveniles^41^ and accelerated blood DNA hypomethylation with age in this cohort^42^, linking rejection history with long-term stress, negative health outcomes, and epigenetic modifications. Here, we explored whether intergenerational transmission of effects occurs and whether such transmission is accompanied by behavioural and molecular alterations. We: (1) analysed demographic and health records to compare lifespan and health outcomes of rejected individuals and their offspring against controls; (2) characterized stress-reactive and affiliative behaviours across two generations; and (3) profiled blood miRNA expression in rejected adults, their well-reared offspring, and controls.

## Results

### Rejected owl monkeys show reduced lifespan and compromised fitness

We first examined whether parental rejection was associated with survival differences using comprehensive demographic records (n=1,372 individuals: 135 rejected, 1,237 non-rejected). Rejected individuals showed significantly reduced lifespan compared to controls (mean age at death: 3.41 years [SD=4.58] vs. 7.03 years [SD=6.02]; Welch’s t-test: p<0.001; Figure 1a). This represents a 51% reduction in lifespan. Among individuals who survived to reproductive age, there was no significant sex difference in lifespan for rejected males versus females (9.05 years [SD=5.55] vs. 8.24 years [SD=5.37]).

**Figure 1.**
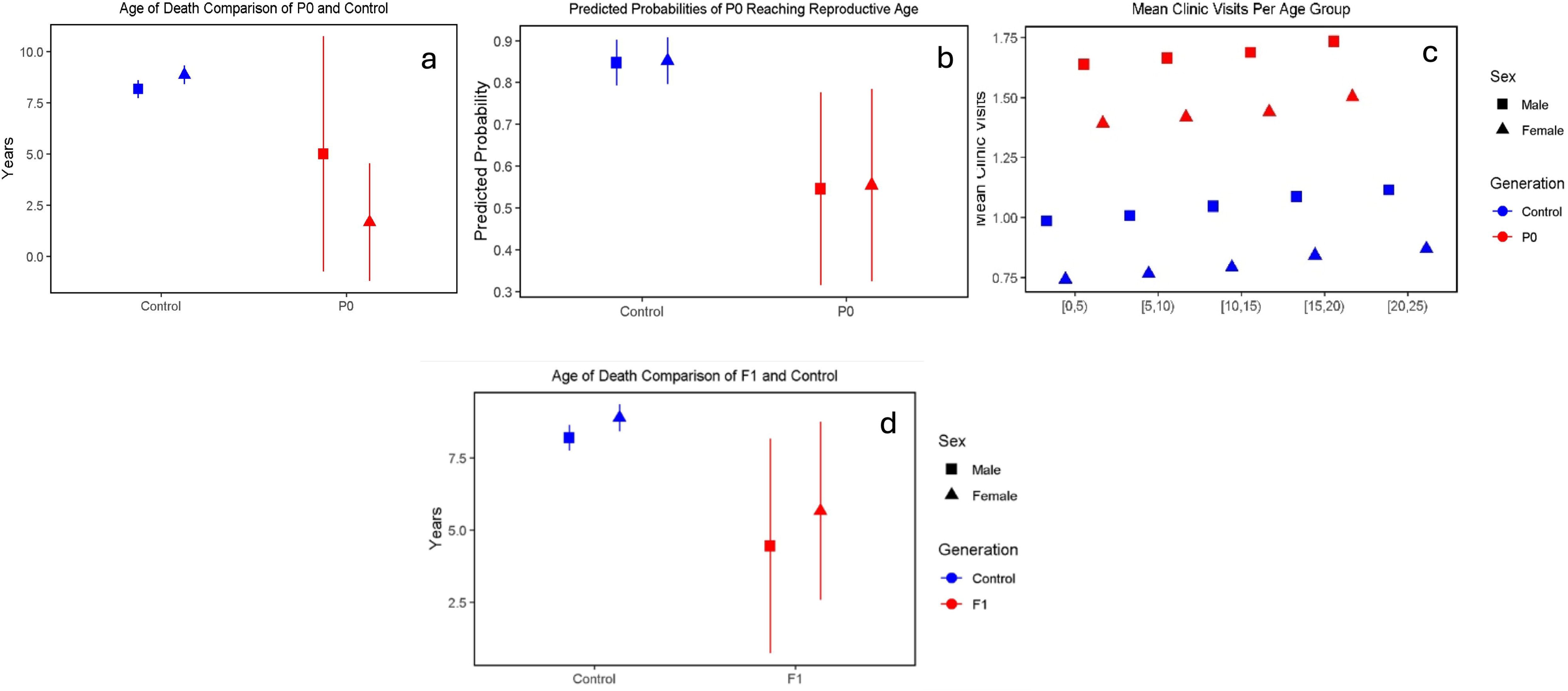
Reduced life expectancy and health in rejected owl monkeys and descendants. (a)Age at death was significantly lower for rejected compared to control monkeys (PO). Rejected individuals (n=135) showed mean age at death of 3.41 years (SD=4.58) compared to 7.03 years (SD=6.02) for controls (n=1,237), representing a 51% reduction in lifespan (Welch’s t-test, p<0.001). (b) Probability of rejected owl monkeys reaching reproductive age (2::1.5 years) was significantly lower than controls (PO). Rejected individuals showed 41.9% reduced odds of reaching reproductive age compared to controls (x^2^ test with Yates’ correction, p<0.01; n=l,161 total: 60 rejected reached reproductive age, 1,101 controls). (c) Clinic visits controlling for age. Rejected individuals (n=6) averaged 1.13 more clinic visits throughout life than controls (n= 53), linear regression, p=0.05. Males and females shown separately, with no significant sex-related differences. (d) Age at death for offspring of rejected parents (n=9) versus controls (n=727), showing 54% reduction in lifespan (mean 3.89 years vs 8.50 years; Welch’s t-test, p=0.018). Males and females shown separately, with no significant sex-related differences. Error bars represent standard deviations. Blue=controls, red=rejected or offspring of rejected.

We next assessed whether rejection affected the probability of reaching reproductive age (defined as ≥1.5 years based on captive breeding data^38^). Among 1,161 individuals analysed, 60 of 152 rejected individuals (39.5%) reached reproductive age compared to 1,101 of 1,237 controls (89.0%). Rejected individuals had significantly lower odds of reaching reproductive age (χ² test with Yates’ correction: p<0.01; odds ratio=0.581; Figure 1b), representing a 41.9% reduction in probability compared to controls.

Given our previous findings of elevated clinic visits in rejected juveniles^42^, we examined whether rejection history predicted health outcomes across the lifespan. Using clinic visits as a proxy for morbidity and controlling for age and sex, we compared rejected and controls visits to clinics (n=59 individuals with full clinic records n=3 female rejected, n=34 non rejected, n=3 males rejected, n=19 non rejected). This preliminary analysis showed that rejected individuals visited clinics 1.13 times more frequently throughout life, p=0.05 (linear regression: β=1.13, SE=0.55, t=2.06, df=55, p=0.044, n=59; Figure 1c). Medical records indicated these visits were primarily for treatment with broad-spectrum antibiotics, suggesting greater susceptibility to infections, a leading cause of mortality in this colony^40^.

### Offspring of rejected parents show reduced survival despite normal care

We next asked whether parental rejection affected offspring fitness, even when offspring themselves received normal parental care. Among 813 individuals with known parental rejection history, offspring of rejected parents (n=31) showed significantly reduced odds of reaching reproductive age compared to offspring of control parents (n=782; generalised logistic regression: β=−0.545, SE=0.156, z=−3.49, p<0.001; odds ratio=0.553; Figure 1d). This represents a 44.7% reduction in probability, similar to the effect of rejection on the directly rejected individuals themselves. Effects were independent of parent sex (β=0.182, SE=0.157, z=1.16, p=0.246) or offspring sex (β=−0.089, SE=0.157, z=−0.57, p=0.571).

Examining lifespan directly in offspring (n=727 controls, n=9 offspring of rejected), we found offspring of rejected parents showed significantly reduced survival (Welch’s t-test: t=2.94, df=9.36, p=0.018; Figure 1d). Mean lifespan was 3.89 years (SD=4.68) for offspring of rejected versus 8.50 years (SD=4.37) for controls, a difference of 4.61 years (95% CI: 1.01–8.21 years), representing a 54% reduction. These results demonstrate that rejection effects on fitness are transmitted intergenerationally, with profound consequences for descendants despite those descendants receiving normal parental care.

Since mechanisms underlying this transmission remain to be fully elucidated and likely involve multiple pathways operating in concert, we explore possible contributors through behavioural and molecular analyses below.

### Rejected adults display altered stress reactivity and reduced parental care

The results of adjusted logistic regression with Bayesian estimation showed that behaviours such as fear (+5%, p=0.03), hugging (+2%, p=0.01), approach (+20%, p=0.01) and hitting others (+10%, p=0.01) increased in parents that were rejected in early life (n=11), whilst hiding on the perch (−44%, p<0.01), carrying their offspring (−7%, p=0.04) and jumping locomotion (−5%, p<0.01) decreased compared to control parents (n=27) (Supplementary File 8). When comparing behavioural differences adjusted for parent sex (Supplementary files 9,10), some behaviours were more evident in females. Mothers with a history of parental rejection showed greater fear (+4%, p=0.01), hugging (+2%, p<0.01), approach (+19%, p=0.07) and agonistic social behaviours such as hitting other monkeys (+11%, p=0.07) and biting (+13%, p=0.30), whilst freezing (−10%, p=0.01), jumping locomotion (−5%, p=0.01) and furtive glancing (−1%, p<0.01) decreased compared to controls. Similarly, fathers with parental deprivation showed increased fear (+5%, p=0.70), approach (+28%, p=0.05) and hitting others (+8%, p=0.07), whilst time spent hiding on the perch (−39%, p=0.01) and hiding behind (−11%, p<0.01) decreased. They also showed decreased baby carrying (−8%, p=0.03) and jumping locomotion (−5%, p<0.01) compared to controls (Figure 2). Only one adult male monkey displayed stereotyped behaviour (head twirl).

**Figure 2.**
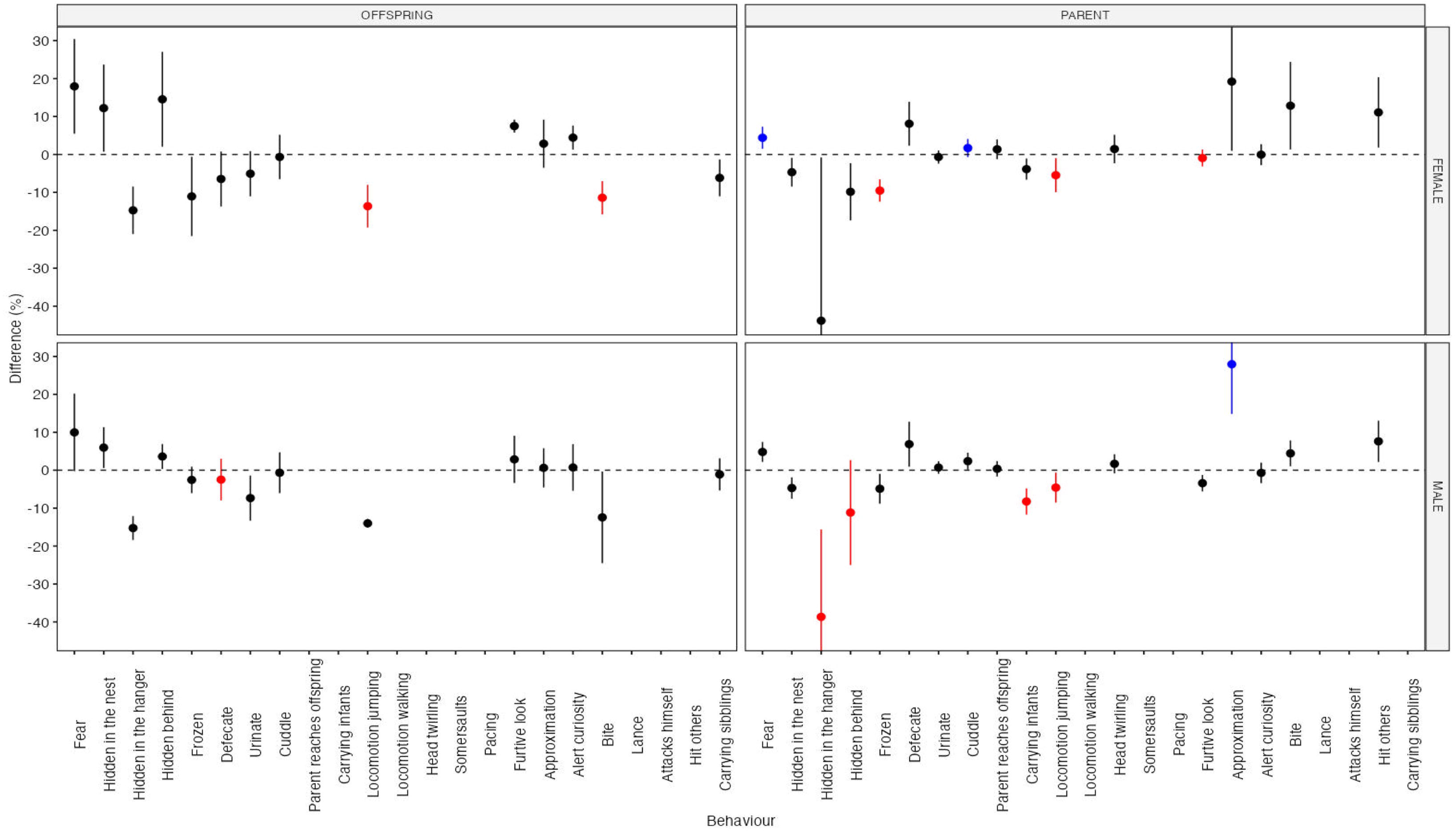
Behavioural differences in rejected parents and offspring compared to controls. Forest plots showing behavioural differences (adjusted percentages controlling for family, sex, and age using Bayesian logistic regression) between rejected and control groups. Upper panels: fathers (n=20 total) and mothers (n=l8 total) (PO). Lower panels: sons (n=lO total) and daughters (n=17 total) (Fl). Black points indicate non-significant differences (p>0.05); blue points indicate significantly higher behaviour frequency in rejected groups (p<0.05); red points indicate significantly lower frequency in rejected groups (p<0.05). Error bars show 95% credible intervals. Rejected adults showed increased fear (+5%, p=0.03), approach/boldness (+20%, p=0.01), and aggression/hitting others (+10%, p=0.01), with decreased hiding on perch (−44%, p<0.01), carrying offspring (−7%, p=0.04), and jumping locomotion (−5%, p<0.01) compared to controls. Offspring of rejected parents showed decreased hiding on perch (−15%, p=0.05), jumping locomotion (−14%, p<0.01), and biting (−12%, p=0.01) compared to offspring of controls. Complete behavioural data provided in Supplementary Files 2-4, 8-9.

### Offspring of rejected parents showed more fearfulness and less locomotion similar to their parents

Using the same statistical analysis, we show offspring of rejected parents (n=13) also display behaviours such as fear (+18%, p=0.14) and hiding in the nest and hiding behind another monkey (+12%, p=0.13; +9%, p=0.8 respectively) increased. However, significant results were only detected amongst behaviours which were decreased for offspring of rejected parents (n=13): hiding on the perch (−15%, p=0.05), jumping locomotion (−14%, p<0.01) and biting (−12%, p=0.01). When considering offspring independently, the same pattern emerges. Female offspring of parents with parental deprivation showed more fear (+18%, p=0.2), hiding in the nest (+12%, p=0.3) and furtive glancing (+7%, p=0.06) but decreased jumping locomotion (−14%, p<0.01) and biting (−11%, p=0.01) compared to female control offspring. Additionally, male offspring showed increased fear (+10%, p=0.4) and decreased jumping locomotion (−14%, p=0.2) and biting (−12%, p=0.3) compared to male control offspring (Figure 2 and Supplementary File 11). Although some differences in behaviours are not statistically significant, numerically they show larger average treatment effects across all groups.

### Female offspring of rejected parents show altered miRNA expression suggesting potential molecular mechanisms

We conducted a differential expression analysis of whole blood miRNAs in animals with a history of rejection, their offspring, and controls (n=47). This analysis was performed to explore whether microRNAs might be associated with parental rejection history and potentially contribute to intergenerational transmission of effects. Whilst this section focused on commenting on significantly dysregulated miRNAs, we provide a complete list of all analysed miRNAs, including those that showed no significant changes (Supplementary File 7). To identify significantly altered miRNAs across these multiple comparisons, we used LIMMA TREND (Partek flow), which provides stringent false discovery rate control (FDR p<0.05).

Our initial analysis demonstrated that, despite the association between adults with a history of rejection and the dysregulation of many miRNAs in blood, none passed the minimum FDR threshold in either rejected males (Figure 3a) or rejected females (Figure 3b). However, we did observe a down-regulated miRNA in males and females (mml-miR-182) with marginal significance (FDR p between 0.05 and 0.10). Subsequently, we compared miRNA in well-reared offspring (F1) of parents who experienced rejection (n=8) with offspring of controls (n=7). This comparison revealed several differentially regulated miRNAs. Again, a miRNA showed common upregulation in female and male offspring (mml-miR-320a) but the effect did not reach significance (FDR p between 0.05 and 0.10).

**Figure 3.**
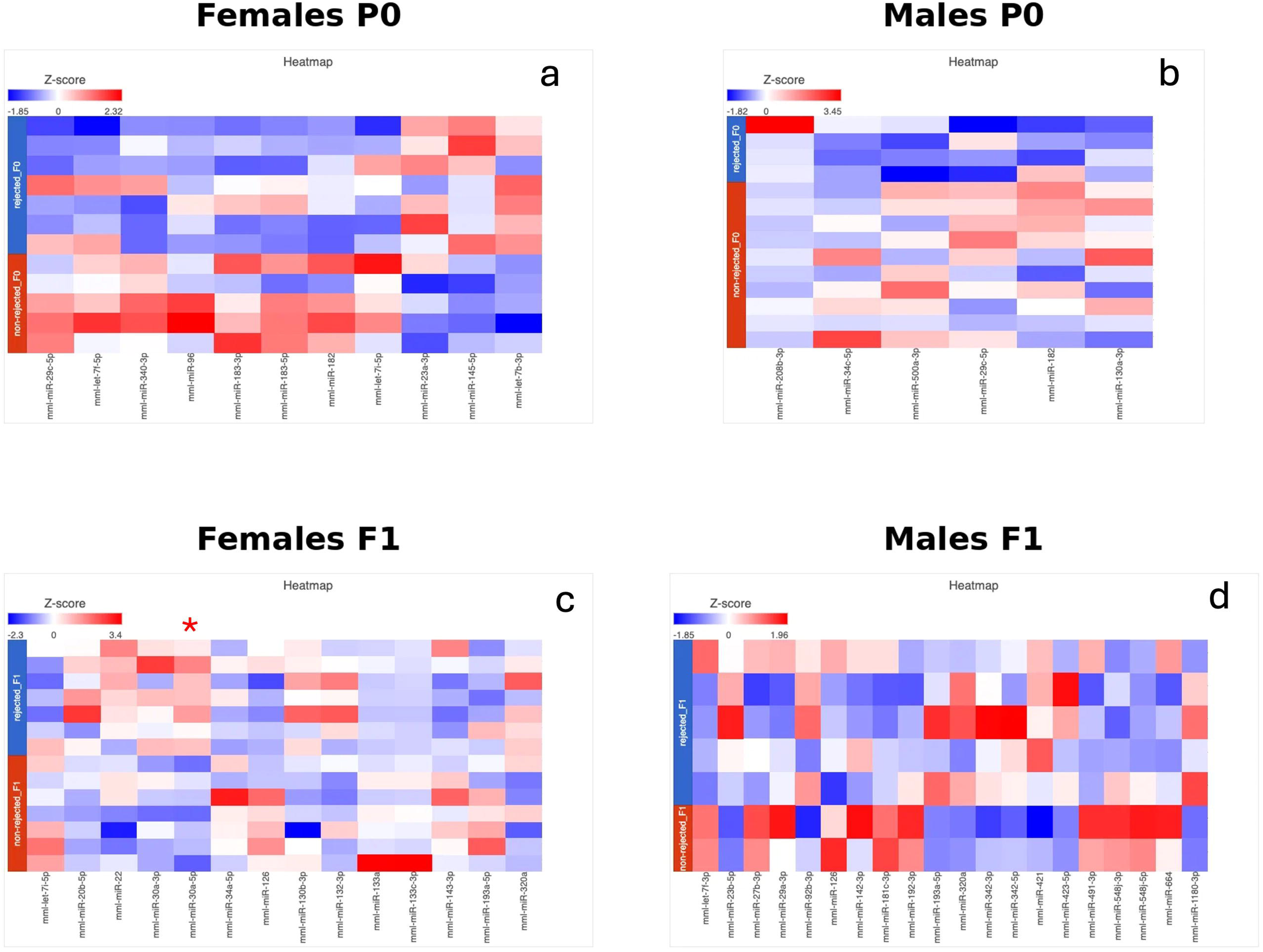
Effects of rejection on miRNA expression in whole blood across two generations of owl monkeys. Heatmaps showing z-score normalised miRNA expression (blue=downregulation, red=upregulation; colour intensity indicates magnitude of dysregulation). Blood samples of 47 were included in analysis after quality control. (a) Rejected females versus control females (PO). (b) Rejected males versus control males (PO). No miRNAs reached significance (FDR p<0.05) in rejected adults; mml-miR-182 showed marginal downregulation (FDR p 0.05-0.10). (c) Well-reared female offspring of rejected parents versus offspring of controls. mml-miR-30a-5p was significantly upregulated specifically in female offspring of rejected parents (FDR p=9.91xI0-^3^, FC=2.09, indicated by red asterisk)-the only miRNA passing stringent multiple testing correction. (d) Well-reared male offspring of rejected parents versus offspring of controls. No miRNAs reached significance. Differential expression analysis used LIMMA TREND with Benjamini-Hochberg FDR correction (Partek Flow). Complete miRNA expression data provided in Supplementary File 7.

However, we did detect significant upregulation of one miRNA: mml-miR-30a-5p, which was significantly overexpressed (FDR p=9.91×10³, FC=2.09) in well-reared female daughters (Figure 3 c,d). This miRNA was the only one to pass stringent multiple testing correction across all comparisons. Among other upregulated miRNAs (p<0.05), we observed the partner miRNA mml-miR-30a-3p. The upregulation of miR-30a-5p specifically in daughters of rejected parents is notable given existing evidence linking this miRNA to stress-related outcomes. miR-30a-5p belongs to a miRNA family implicated in mediating social stress and behavioural disorders in mammals^43^. In humans, miR-30a-5p is upregulated in blood plasma of individuals with depression who experienced early-life adversity and in brains of rats exposed to early life stress^44^.

## Discussion

### Parental rejection predicts reduced lifespan and compromised fitness

The 51% reduction in lifespan observed in rejected owl monkeys, despite veterinary care and identical post-weaning conditions as controls, demonstrates that early-life stress has profound, lasting effects on mortality even when environmental factors are controlled. Our finding that rejection remains a significant predictor of morbidity throughout life, with rejected individuals requiring 1.13 times more clinic visits, suggests chronic health vulnerability may contribute to reduced longevity.

The predominant reason for clinic visits was antibiotic treatment, suggesting increased susceptibility to infections. Given that pneumonia and enteritis are among leading causes of mortality in this colony^40^, heightened infection risk may account for increased morbidity, paralleling observations in rhesus macaques^9^. Stress-induced immune dysregulation is well-documented in ELA research^8^. As we previously observed HPA axis alterations in rejected individuals, results from our present study could reflect compromised immune function.

High incidences of nephropathies and cardiomyopathies are documented in this colony^40,45,46^ and are associated with stress exposure and sudden death in owl monkeys. Whether rejected individuals show higher prevalence of these chronic conditions remains to be tested. The exact origins of reduced survival in rejected owl monkeys warrant further investigation, but the combination of increased infection susceptibility and potential stress-related pathologies likely contributes to the observed mortality patterns.

Intergenerational transmission of reduced fitness despite normal parenting. The most striking finding is that offspring of rejected parents, despite receiving normal parental care themselves, showed nearly identical reductions in survival to reproductive age (44.7% reduction) and lifespan (54% reduction) as their rejected parents. This demonstrates genuine intergenerational transmission rather than simply persistent effects of poor rearing.

These results extend findings in wild baboons^30^ to a controlled captive setting, demonstrating that intergenerational transmission occurs independent of environmental variation. All sources of life history variation typical in wild populations (resource availability, predation, disease exposure) are controlled in captivity. All primates live in identical conditions, and juveniles are moved to new enclosures by management staff at the age of weaning. The persistent survival deficit in offspring of rejected parents therefore cannot be attributed to external environmental factors or differences in parental investment such as resource competition observed in the wild.

Retrospective analysis of health outcomes in F1 offspring to explain earlier mortality was not possible, as medical records in this colony were created only when individuals visited clinics, and most F1 individuals recorded for our study died before our study began. This may indicate that negative health outcomes were subclinical and went undetected.

The persistence of reduced survival across generations, despite controlled environmental conditions and seemingly equal parental care, suggests that biological embedding of parental stress is transmitted. Our behavioural and molecular analyses below explore potential pathways underlying this transmission.

### Behavioural transmission

altered stress reactivity, boldness, and parental care. Rejected adults showed a distinct behavioural profile characterised by heightened fearfulness yet paradoxically increased boldness (approach behaviour) and reduced hiding. This combination has been observed in rodents following maternal separation^47,48^ and may reflect dysregulated threat assessment or risk-taking. Increased emotionality was also detected in rhesus macaques whose fathers experienced early-life separation^29^. Offspring of rejected parents showed similar trends towards increased fearfulness and boldness, though these did not reach statistical significance, potentially reflecting subtle intergenerational transmission detectable with larger samples.

The finding that rejected fathers showed significantly reduced infant carrying is particularly striking. Owl monkey fathers are renowned for intensive paternal care, carrying infants up to 90% of the time^18,38,39,49^, a trait considered an evolutionary specialisation. Our results demonstrate unexpected plasticity in this supposedly constrained behaviour, suggesting rejection experience can fundamentally alter social bonding and caregiving motivation. The origins of this caregiving deficit are unknown but could stem from disrupted attachment formation during early development^50^. This parallels findings in rhesus macaques where maternal rejection predicts abusive parenting in the next generation^27^, though our study extends effects to paternal care.

Offspring of rejected parents showed intergenerational transmission of reduced locomotion in response to stress. Decreased mobility has been reported in ELA-exposed rhesus macaques^51^ and marmosets^52^ and is thought to reflect motor system inhibition mediated by fronto-amygdala circuits-a potential stress coping mechanism^53^. The consistency of this response across generations suggests heritable alterations in stress-responsive neural circuits. Reduced mobility in stress contexts may result from HPA axis overactivity^54^.

Interestingly, biting behaviour showed opposite patterns across generations: rejected adults showed increased biting, whilst their offspring showed decreased biting. As biting is part of the maternal behavioural repertoire during normal weaning in owl monkeys^55^, elevated biting in rejected parents could contribute to perpetuating cycles of rejection, observed in other non-human primates exposed to maternal neglect^27^. However, the suppression of this response in studied offspring, despite their elevated emotionality, suggests adaptive regulation of aggressive responses, potentially representing positive stress coping in well-reared descendants.

Altered behaviours cannot be explained solely by learning or insufficient developmental experience. Rejected parents can successfully rear some offspring whilst rejecting others, indicating rejection is not deterministic. Moreover, offspring showed behavioural patterns opposite to their parents (e.g., decreased biting), arguing against simple social learning. Together, our findings indicate intergenerational transmission of behavioural plasticity associated with rejection experience. This is ecologically relevant given that owl monkey fathers provide extensive care both in captivity^18,38,40,55^ and in the wild^39,49^, and all owl monkey mothers reject their babies to pass on caring duties to fathers during natural weaning.

### Altered miRNA expression in offspring

preliminary evidence for molecular mechanisms. The significant upregulation of mml-miR-30a-5p specifically in female offspring of rejected parents provides preliminary evidence that miRNAs may contribute to intergenerational stress transmission, though functional significance remains to be established. miR-30a-5p belongs to a miRNA family implicated in mediating social stress and behavioural disorders in mammals^43^. In adult humans with depression and early-life adversity history, miR-30a-5p is upregulated in blood plasma^44^, and the same study demonstrated altered DNA methylation patterns at this locus in brains of rats exposed to ELA, demonstrating the association between this circulatory biomarker and brain alterations in response to ELA. The functional significance of miR-30a-5p overexpression in owl monkeys is unknown. If functional, miR-30a-5p dysregulation could increase vulnerability to stress disorders in owl monkeys; elevated expression in daughters of rejected parents could contribute to perpetuating alterations in this population, particularly given females determine lactation, early offspring care, and initiate weaning rejection. However, miR-30a-5p dysregulation could also indicate other underlying physiological differences in blood between these cohorts. The very limited number of altered miRNAs despite profound phenotypic differences in rejected individuals and their offspring suggests multiple mechanisms are at play. At a genomic level, we have also observed accelerated DNA hypomethylation in rejected owl monkeys^42^. Altogether, our study in rejected owl monkeys supports the view that epigenome-wide effects are associated with ELA^28,33^ and that miRNAs are associated with transmission of altered phenotypes.

### Limitations

Several important limitations should be acknowledged. Our study is observational and correlational. Whilst the large demographic sample (n=1,372) provides robust evidence that intergenerational effects occur, we cannot experimentally manipulate parental rejection or control all potential confounding factors.

Could rejection itself have a heritable genetic component that also predisposes to poor health outcomes? Several observations argue against this as the sole explanation: rejected individuals successfully reared some offspring whilst rejecting others. Quality of the rejected owl monkeys themselves is a possible cause, as proposed in rejection events in humans and non-human primates^56,57^, and could explain differences in naïve female offspring. However, some rejected individuals, and their offspring enjoyed long healthy lives and reproduced successfully. Anecdotal reports from veterinary inspections showed no apparent differences, paralleling other observations of rejection in New World primates^58^. Sources of individual variability such as genetic or chromosomal aberrations that might predispose individuals to rejection or to suffer from intergenerational effects of rejection will be analysed in future studies.

Further, we could not assess prenatal conditions, maternal health during pregnancy, or subtle early postnatal experiences that might differ between rejected and control families. Whilst all families lived in identical conditions post-weaning, earlier developmental differences could contribute to observed effects. Some behavioural differences showed non-significant trends; larger studies might clarify which changes are robust.

Future work should address several critical questions raised by our findings. First, whilst we demonstrate that rejection harms offspring fitness, the evolutionary persistence of rejection behaviour remains paradoxical. Investigating whether rejecting behaviour itself confers fitness benefits to parents-such as extended lifespan or enhanced reproductive output-could resolve this apparent contradiction and clarify whether rejection represents an adaptive maternal strategy under captivity. Second, the molecular pathways underlying the negative effects on rejected offspring warrant investigation. Third, characterising rejection behaviour transmission itself-determining whether rejected individuals are more likely to reject their own offspring and identifying associated molecular signatures-would establish whether rejection propensity is epigenetically inherited alongside the physiological and behavioural alterations we observed. Finally, extending molecular analyses to germline tissues, conducting prospective longitudinal studies monitoring maternal physiology during pregnancy, and examining F2 generation outcomes would clarify timing and mechanisms of epigenetic transmission.

### Broader evolutionary and biomedical significance

From an evolutionary perspective, intergenerational transmission of stress effects may represent predictive adaptive responses, where parental experiences inform offspring phenotypes in preparation for similar environments^1,2^. However, whether the effects we observe represent adaptive calibration or simply detrimental consequences of stress exposure remains unclear. In captivity, environmental conditions differ radically from wild habitats, potentially rendering responses that were once adaptive into maladaptive outcomes. The mixed pattern of behavioural changes in offspring, including both potentially detrimental effects (reduced locomotion, heightened fear) and possible adaptive stress coping (reduced aggression), suggests complex interactions between inherited vulnerability and developmental plasticity rather than simple pathology.

For biomedical research, our findings underscore that early-life history shapes lifelong health trajectories and that parental history profoundly influences offspring outcomes even when offspring themselves receive apparently normal care. This has implications for the developmental origins of health and disease (DOHaD) framework^1^, highlighting that intergenerational effects may confound interpretation of results from primate biomedical models where parental early-life history is often unknown or uncontrolled. Practical implications. Our findings have important practical implications for captive primate management. Complete parental rejection affects approximately 5-10% of births in this owl monkey colony and has been documented in other owl monkey colonies^59^ and across primate taxa^13–26^. Current practice is mixed, but in some facilities, rejected infants are hand-reared and then returned to breeding populations. Our results suggest this approach may inadvertently propagate intergenerational effects, compromising welfare and fitness of subsequent generations. Breeding programmes should consider enhanced monitoring of descendants of rejected individuals for health issues, careful documentation of rejection history in pedigree databases, and investigation of environmental or management factors that may reduce rejection rates. These considerations are important for colonies producing animals for biomedical research, where behavioural and physiological consistency is critical.

## Conclusion

We identified robust intergenerational effects of parental rejection in owl monkeys, manifesting as reduced survival and altered stress and social behaviours transmitted across generations despite normal parental care of offspring. Blood miRNA analysis revealed sex-specific dysregulation in female offspring, providing preliminary evidence that epigenetic mechanisms are associated with transmission, though multiple pathways likely operate. The central finding-that parental rejection effects are transmitted intergenerationally, reducing offspring survival by 54% despite normal care-is robustly supported by large-scale demographic data. The mechanisms underlying this transmission remain incompletely understood and warrant further investigation. This study represents the first comprehensive characterisation of intergenerational rejection effects in a captive non-traditional primate model and has important implications for primate welfare, captive breeding practices, and understanding intergenerational transmission of stress effects relevant to human health.

## Materials and Methods

### Ethics statement

All experimental procedures adhered to the American Society of Primatologists (ASP) Principles for the Ethical Treatment of Non-Human Primates. The study was approved by the Welfare and Research Committee of San Marcos University and the UNMSM ethics committee. Research permits were issued by the Peruvian authority SERFOR to Dr. Paredes (AUT-IFS-2021-040 and RDG NO D000334-2021-MIDAGRI-SERFOR-DGGSPFFS).

### Study location and population

Research was conducted at the Centre for Reproduction and Conservation of Non-Human Primates (CRCP) of IVITA at Universidad Nacional Mayor de San Marcos (UNMSM), Iquitos, Peru. The colony has maintained detailed demographic and medical records since establishment. Rejection events (defined as maternal cessation of lactation and/or paternal aggression towards infants) are documented, and rejected infants are rescued via nursery-rearing.

### Housing and husbandry

Families were housed in single enclosures (2 m³: 1 m width × 1 m depth × 2 m height) with natural photoperiod. Owl monkeys received daily feeding of seasonal fruits, balanced dry food, and water ad libitum.

### Sample composition

Sample sizes varied by analysis due to data availability (Supplementary File 1). For survival analysis, we included 1,372 individuals (135 rejected, 1,237 control). Reproductive age analysis included 1,161 individuals (60 rejected who reached reproductive age, 1,101 controls). Analysis of intergenerational effects on reproductive age included 813 individuals (31 offspring of rejected parents, 782 offspring of controls). Intergenerational effects on lifespan were analysed in 736 individuals (9 offspring of rejected, 727 controls). Behavioural observations were conducted on 65 individuals across 10 families spanning 2 generations. MicroRNA analysis initially included 48 samples, reduced to 47 after quality control, from 10 families across 2 generations selected for behavioural study.

### Family cohort selection

For behavioural and molecular studies, we identified 10 families including at least one adult with documented rejection history who survived to adulthood and had offspring at the time of study. Age-matched control families with offspring were selected. The rejected group (P0) comprised 11 rejected adults (6 females, 5 males), 8 well-reared breeding partners (3 females, 5 males), and 14 juvenile offspring (F1: 9 females, 5 males). The control group (P0) included 19 well-reared adults (9 females, 10 males) and 13 juvenile offspring (F1: 8 females, 5 males). Selection criteria ensured groups were matched for age, housing conditions, and offspring number and age.

### Demographic and health analyses

We compared age at death between rejected and control individuals using Welch’s two-sample t-test. For individuals alive at data collection, current age was used as censored data. Mean age at “death or data collection” was calculated with standard deviations. Figures were generated in R.

Based on captive breeding data^38^, reproductive age was defined as ≥1.5 years. Pearson’s chi-square test with Yates’ continuity correction assessed associations between rejection status and probability of reaching reproductive age.

Generalised logistic regression examined whether parental rejection predicted offspring reaching reproductive age, controlling for parent sex and offspring sex. Welch’s two-sample t-test compared lifespan between offspring of rejected parents and controls.

Linear regression assessed whether rejection history predicted clinic visits throughout life, controlling for age and sex.

### Behavioural characterisation

We conducted a modified human-intruder test^24^ one month before blood sampling. Testing occurred 1900-2000 hours when monkeys are most active^60^. Monkeys were exposed to low-intensity stressors: light stimulus (camera light), human presence (unfamiliar observer, “intruder”), and mechanical stimulus (cage door opening).

Three one-minute observation sessions were video-recorded per individual with five-minute inter-session intervals. Behaviours were quantified using ethograms developed from previous studies in *Aotus* (Supplementary File 2). Frequency and duration of 18 behaviours were recorded, including stress responses (fear, freezing, hiding), locomotion, social behaviours (grooming, carrying offspring), and agonistic behaviours (biting, hitting). Raw behavioural percentages by sex and generation are provided in Supplementary File 3.

Behavioural events were observed using Behavioural Observation Research Interactive Software (BORIS). Total duration and number of episodes per individual were quantified. Behaviours were evaluated by two independent observers to avoid biased observations, with inter-rater reliability ≥95%. Complete behavioural data are provided in Supplementary File 4.

### Statistical analysis of behaviour

We used Bayesian logistic regression to estimate behavioural differences, adjusting for family/cage effects, sex, and age category. Parental rejection history was the main predictor. The outcome was binary (0/1) indicating whether an individual displayed each specific behaviour at least once during observation periods (0 = no behaviour observed, 1 = at least one event observed).

Predicted/adjusted frequencies were compared using exact Wilcoxon rank-sum tests. Average treatment effects were estimated using G-computation, assigning all individuals to each treatment condition and estimating differences between predicted posterior distributions. Statistical models and priors are described in Supplementary File 5. Results with p<0.05 were considered significant.

### Blood collection

Blood samples were collected from a subset of 48 owl monkeys (rejected P0: 8 females, 4 males; offspring of rejected F1: 7 females, 5 males; control P0: 10 females, 8 males; offspring of controls F1: 7 females, 2 males) during routine veterinary procedures under ketamine sedation (8 mg/kg). Samples were obtained via femoral venipuncture using 3 ml syringes with 1“×23G needles into EDTA Vacutainer tubes. From each sample, 0.5 ml was spotted onto filter paper and air-dried at room temperature.

### MicroRNA extraction and sequencing

Total RNA was extracted from three 6-mm punches per dried blood spot using the mirVana microRNA Isolation Kit (Thermo Fisher Scientific, AM1560) following manufacturer’s protocols^61,62^. Briefly, spots were suspended in cell disruption buffer, denaturing solution was added, followed by acid phenol:chloroform extraction. After centrifugation at 4°C, the aqueous phase was collected, mixed with 100% ethanol, and applied to filter cartridges. After washing, RNA was eluted in pre-heated elution buffer.

Total RNA was concentrated using the RNeasy MinElute Cleanup Kit (Qiagen, 74204). Lysis buffer containing guanidine thiocyanate was added to RNA samples, followed by ethanol. The mixture was applied to RNeasy MinElute spin columns, washed, and high-quality concentrated RNA was eluted in 12 μl RNase-free water and stored at −80°C.

Libraries were prepared using QIAseq miRNA Library Kit (Qiagen) according to manufacturer’s instructions. Library quality was assessed using NanoDrop 8000 (Thermo) and TapeStation 4200 (Agilent) before sequencing on NextSeq 2000 (Illumina Inc.).

### Bioinformatic preprocessing and differential expression analysis

Raw sequencing reads were trimmed using Trim Galore v0.6.6 to remove adapters and low-quality bases. Trimmed reads were aligned to the owl monkey genome (*Aotus nancymaae*, Anan_2.0) using STAR v2.7.3a. Gene-level counts were generated using HTSeq-count.

Raw counts were loaded into Partek Flow (https://www.partek.com/partek-flow/) for analysis. Counts were normalised (counts per million, CPM), filtered to remove unexpressed genes, and principal component analysis (PCA) was performed to evaluate sample variability and detect outliers. One sample (T-1499, P0 female control) was excluded as an outlier based on PCA (Supplementary File 6). After filtering, approximately 3,000 sequences matched the owl monkey genome.

Since the *Aotus nancymaae* genome is not fully annotated for miRNAs, we mapped owl monkey small non-coding RNA sequences to the rhesus macaque (*Macaca mulatta*) genome (Mmul_10, GCA_003339765.3) to identify miRNA homologues.

Normalised expression ratios were analysed in Partek Flow. We used the LIMMA TREND method to detect differentially expressed miRNAs between rejected adults versus control adults (stratified by sex) and offspring of rejected parents versus offspring of controls (stratified by sex). Multiple testing correction was performed using the Benjamini-Hochberg procedure to control false discovery rate (FDR). Significance thresholds were set at FDR p<0.05 and fold change ≥1.5 or ≤−1.5. All reported p-values are FDR-adjusted unless stated otherwise. Complete miRNA expression data including non-significant results are provided in Supplementary File 7.

For heatmap visualisation, miRNA expression data were log-transformed and z-scores were calculated by subtracting the mean expression of each miRNA across all samples and dividing by the standard deviation, allowing standardised comparison across multiple miRNAs.

### Statistical software and reproducibility

All statistical analyses were conducted using R version 4.1.0 for demographic and survival analyses, RStan for Bayesian behavioural analyses, Partek Flow version 10.0 for miRNA differential expression, and BORIS version 7.9.18 for behavioural coding.

### Power and sample size considerations

Sample sizes were determined by availability of individuals meeting inclusion criteria in this colony rather than prospective power calculations. For demographic analyses, large sample sizes (n=1,389) provided excellent power (>95%) to detect moderate effect sizes (Cohen’s d ≥0.3). For behavioural and miRNA analyses, smaller sample sizes (n=47-65) limited power to detect small effects but provided adequate power (80%) to detect moderate-to-large effects (Cohen’s d ≥0.7 for behaviour; fold change ≥1.5 for miRNA with FDR correction).

### Blinding

Behavioural observers were not fully blinded to rejection status, as this information was part of colony records. However, observers were not informed of specific study hypotheses regarding which behaviours were expected to differ, and coding was performed by two independent observers to minimise bias. For miRNA analysis, laboratory personnel processing samples and bioinformaticians conducting analyses were blinded to sample identity and rejection status.

## Data availability

All data supporting the findings of this study are available within the manuscript and supplementary files. Raw miRNA sequencing data will be deposited in NCBI Gene Expression Omnibus (GEO) upon acceptance. Summary of biological sampling across analysis (de-identified) is provided in Supplementary File 1. Behavioural data are provided in Supplementary Files 2-4, 8-11. miRNA expression data are provided in Supplementary File 7. Analysis code for demographic and behavioural analyses is provided in Supplementary File 5. Additional data supporting the findings are available from the corresponding author upon reasonable request, subject to ethical restrictions to protect animal identities and colony security.

## Author Contributions

Conceptualization: UMP, JF, MC

Data curation: NSP, JF, MC, UMP

Formal analysis: JF, MC, JB, PY, UMP

Funding acquisition: UMP, NSP, PY

Investigation: JF, MC, FOCP

Methodology: JF, MC, JB, CM, PY, UMP

Project administration: UMP, NSP

Resources: UMP, NSP, PY

Software: JF, FOCP, JB

Supervision: UMP, NSP

Validation: JF, MC, JB, PY

Visualization: JF, MC, UMP

Writing - original draft: JF, MC, FOCP, UMP

Writing - review & editing: All authors

## Acknowledgments

We thank the primate care staff and laboratory scientists at IVITA-Iquitos led by Nofre Sanchez, for assisting with data and sample collection. We are grateful to staff at the Blizard Institute Genome centre for assistance with molecular analyses. We thank Prof. Vardhman Rakyan for helpful discussions.

