## Supplementary material 1-11 for "Intergenerational transmission of parental rejection effects on survival, behaviour, and miRNA expression in owl monkeys"

| **Analysis** | **Total N** | **Rejected** | **Non-rejected / Controls** | **Notes** |
| --- | --- | --- | --- | --- |
| **Survival analysis** | 1,372 | 135 | 1,237 | Full colony |
| **Reproductive age analysis** | 1,161 | 60 | 1,101 | Individuals ≥1.5 years old |
| **Intergenerational effects on reproductive age** | 813 | 31 (offspring of rejected parents) | 782 (offspring of controls) | — |
| **Intergenerational effects on lifespan** | 736 | 9 (offspring of rejected parents) | 727 (offspring of controls) | — |
| **Family cohort / behavioural observations** | 65 | 33 adults + offspring | 32 adults + offspring | 65 individuals from 10 families spanning 2 generations; includes F0 adults and F1 offspring with recorded behavioural observations, including rejection-related behaviours |
| **miRNA sequencing** | 47 | 10 adults + 8 offspring | 22 adults + 7 offspring | From 10 families across 2 generations; initially 48 samples, 1removed after QC; includes F0 and F1 individuals with behavioural and parental rejection data. |

**Supplementary File 2. Ethogram used to collect data for all the listed behaviours**

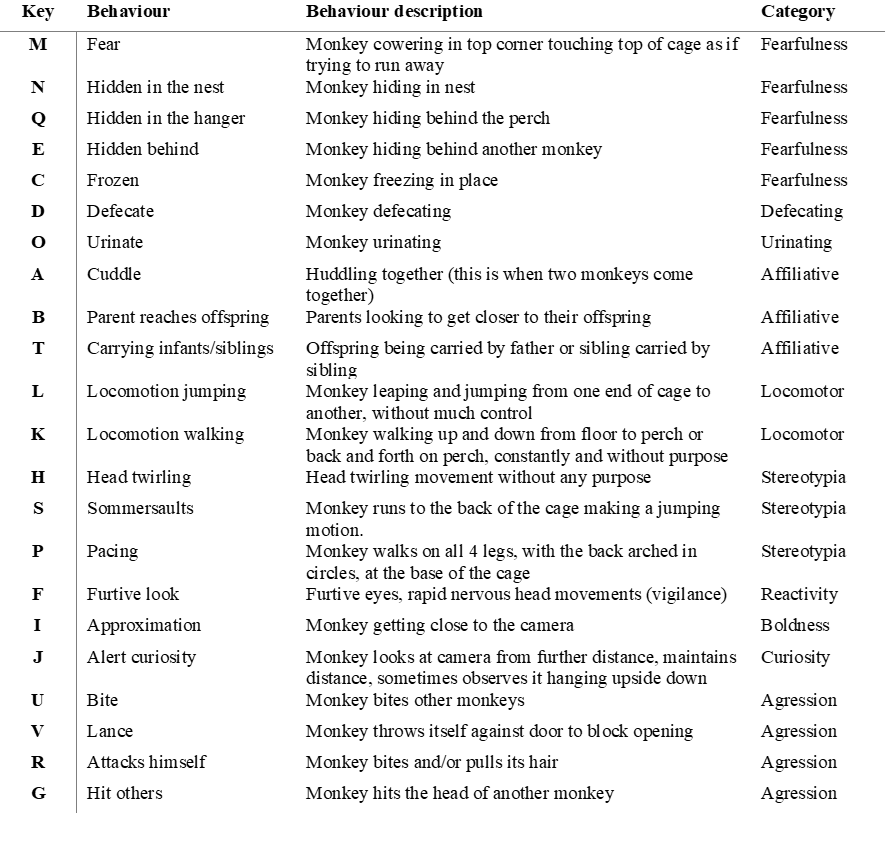

**Supplementary File 3. Unadjusted percentages of each behaviour by sex**

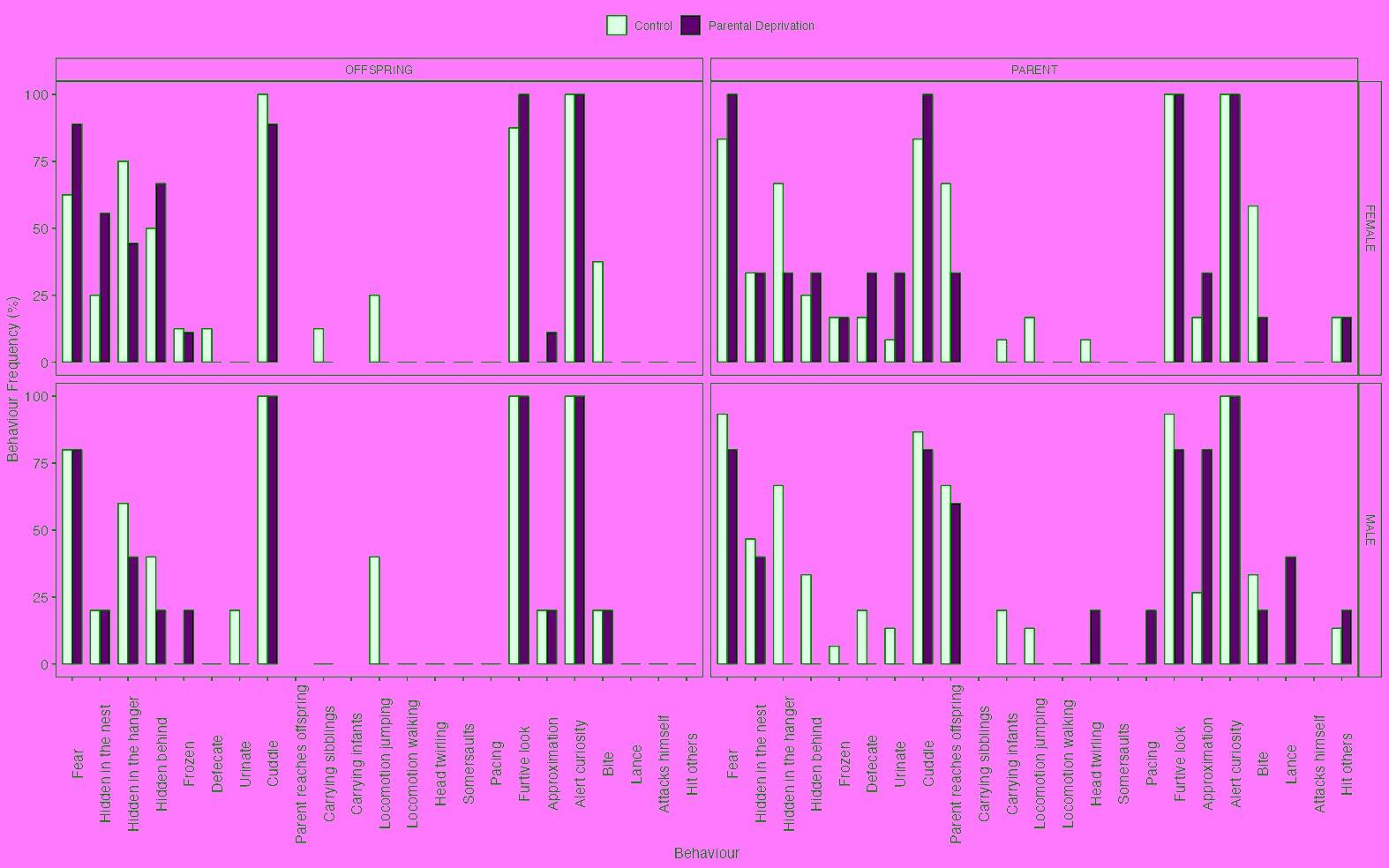

Unadjusted percentages of individuals displaying each behaviour at least once by sex and age categories. Parents with parental rejection ( n=11): 5 males+6 females; Parents without parental rejection (n=27): 15 males+12 females; Offspring from one parent with parental rejection (n=14): 5 males+9 females, and Offspring from parents without parental rejection (n=13): 5 males+9 females.

**Supplementary file 4. Ethogram data (excel file)**

This file is ready and translated to English here: [Suppl File 3 - Ethogram Data.xlsx](https://docs.google.com/spreadsheets/d/1A33DRA_O9CR0T2qhcQGbO78pjTKBqByG/edit?usp=sharing&ouid=103699935812651173190&rtpof=true&sd=true)

**Supplementary file 5. Ethogram data analysis (R file)** [Suppl File 5 - Ethogram data analysis.qmd](https://drive.google.com/file/d/1zhiJllXLJUWP5BkxjiSKHWd-K3rcs1Tp/view?usp=sharing)

**Supplementary file 6.** Identification of Outliers based on PCA analysis of all samples which were RNA-sequenced. Based on distribution we excluded sample T-1499.

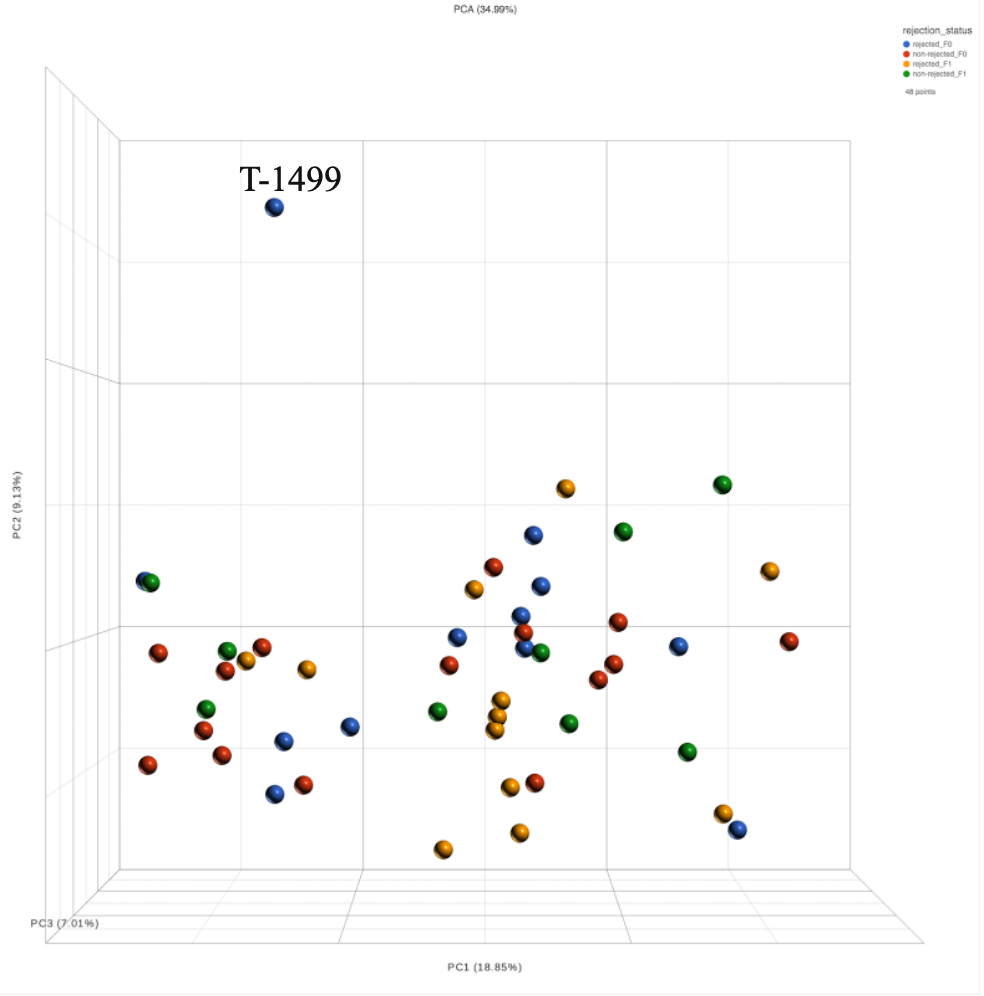

**Supplementary File 7. Summary from miRNA analysis**

| **microRNA ID** | **Log2 FC** | **P-value (p<0.05)** | **FDR adjusted p** | **Comparison** |
| --- | --- | --- | --- | --- |
| **mml-miR-500a-3p** | **-1.56** | **0.00867** | **0.984** | **F0 Male (Rejected vs Non-Rejected)** |
| **mml-miR-34c-5p** | **-2.87** | **0.0158** | **0.984** | **F0 Male (Rejected vs Non-Rejected)** |
| **mml-miR-29c-5p** | **-1.66** | **0.0164** | **0.984** | **F0 Male (Rejected vs Non-Rejected)** |
| **mml-miR-208b-3p** | **3.72** | **0.0166** | **0.984** | **F0 Male (Rejected vs Non-Rejected)** |
| **mml-miR-130a-3p** | **-1.75** | **0.0255** | **0.984** | **F0 Male (Rejected vs Non-Rejected)** |
| **mml-miR-182** | **-1.6** | **0.0282** | **0.984** | **F0 Male (Rejected vs Non-Rejected)** |
| **mml-miR-30a-5p** | **-1.75** | **0.0397** | **0.984** | **F0 Male (Rejected vs Non-Rejected)** |
| **mml-miR-150-3p** | **-2.49** | **0.0467** | **0.984** | **F0 Male (Rejected vs Non-Rejected)** |
| **mml-miR-145-5p** | **2.4** | **0.00807** | **0.702** | **F0 Female (Rejected vs Non-Rejected)** |
| **mml-let-7b-3p** | **1.74** | **0.00961** | **0.702** | **F0 Female (Rejected vs Non-Rejected)** |
| **mml-miR-182** | **-1.86** | **0.0123** | **0.702** | **F0 Female (Rejected vs Non-Rejected)** |
| **mml-miR-96** | **-2.44** | **0.0136** | **0.702** | **F0 Female (Rejected vs Non-Rejected)** |
| **mml-let-7i-5p** | **-1.32** | **0.0153** | **0.702** | **F0 Female (Rejected vs Non-Rejected)** |
| **mml-miR-183-3p** | **-2.27** | **0.0156** | **0.702** | **F0 Female (Rejected vs Non-Rejected)** |
| **mml-miR-23a-3p** | **1.42** | **0.0336** | **0.895** | **F0 Female (Rejected vs Non-Rejected)** |
| **mml-miR-183-5p** | **-1.99** | **0.0386** | **0.895** | **F0 Female (Rejected vs Non-Rejected)** |
| **mml-miR-340-3p** | **-1.66** | **0.0387** | **0.895** | **F0 Female (Rejected vs Non-Rejected)** |
| **mml-miR-7** | **-1.53** | **0.0426** | **0.895** | **F0 Female (Rejected vs Non-Rejected)** |
| **mml-let-7f-5p** | **-1.31** | **0.0458** | **0.895** | **F0 Female (Rejected vs Non-Rejected)** |
| **mml-miR-548j-5p** | **-7.81** | **0.00106** | **0.146** | **F1 Male (from Rejected vs Non-Rejected F0)** |
| **mml-miR-342-3p** | **2.71** | **0.00108** | **0.146** | **F1 Male (from Rejected vs Non-Rejected F0)** |
| **mml-miR-1180-3p** | **3.66** | **0.00681** | **0.452** | **F1 Male (from Rejected vs Non-Rejected F0)** |
| **mml-miR-181c-3p** | **-3.32** | **0.0112** | **0.452** | **F1 Male (from Rejected vs Non-Rejected F0)** |
| **mml-miR-192-3p** | **-1.97** | **0.0114** | **0.452** | **F1 Male (from Rejected vs Non-Rejected F0)** |
| **mml-miR-342-5p** | **2.45** | **0.0117** | **0.452** | **F1 Male (from Rejected vs Non-Rejected F0)** |
| **mml-miR-193a-5p** | **5.74** | **0.0139** | **0.452** | **F1 Male (from Rejected vs Non-Rejected F0)** |
| **mml-miR-421** | **1.66** | **0.014** | **0.452** | **F1 Male (from Rejected vs Non-Rejected F0)** |
| **mml-miR-23b-5p** | **2.49** | **0.015** | **0.452** | **F1 Male (from Rejected vs Non-Rejected F0)** |
| **mml-let-7f-3p** | **-2.12** | **0.0185** | **0.502** | **F1 Male (from Rejected vs Non-Rejected F0)** |
| **mml-miR-92b-3p** | **1.96** | **0.022** | **0.543** | **F1 Male (from Rejected vs Non-Rejected F0)** |
| **mml-miR-491-3p** | **-2.36** | **0.0259** | **0.575** | **F1 Male (from Rejected vs Non-Rejected F0)** |
| **mml-miR-664** | **-1.88** | **0.034** | **0.575** | **F1 Male (from Rejected vs Non-Rejected F0)** |
| **mml-miR-548j-3p** | **-1.67** | **0.0343** | **0.575** | **F1 Male (from Rejected vs Non-Rejected F0)** |
| **mml-miR-142-3p** | **-2.08** | **0.0428** | **0.575** | **F1 Male (from Rejected vs Non-Rejected F0)** |
| **mml-miR-27b-3p** | **-1.63** | **0.043** | **0.575** | **F1 Male (from Rejected vs Non-Rejected F0)** |
| **mml-miR-423-5p** | **2.11** | **0.0448** | **0.575** | **F1 Male (from Rejected vs Non-Rejected F0)** |
| **mml-miR-126** | **-1.81** | **0.0468** | **0.575** | **F1 Male (from Rejected vs Non-Rejected F0)** |
| **mml-miR-29a-3p** | **-1.81** | **0.047** | **0.575** | **F1 Male (from Rejected vs Non-Rejected F0)** |
| **mml-miR-320a** | **1.65** | **0.0486** | **0.575** | **F1 Male (from Rejected vs Non-Rejected F0)** |
| **mml-miR-30a-5p*** | **2.18** | **0.0000366** | **0.00991** | **F1 Female (from Rejected vs Non-Rejected F0)** |
| **mml-miR-143-3p** | **-1.82** | **0.0137** | **0.751** | **F1 Female (from Rejected vs Non-Rejected F0)** |
| **mml-miR-193a-5p** | **-2.2** | **0.018** | **0.751** | **F1 Female (from Rejected vs Non-Rejected F0)** |
| **mml-miR-22** | **1.49** | **0.0185** | **0.751** | **F1 Female (from Rejected vs Non-Rejected F0)** |
| **mml-miR-133a** | **-4.12** | **0.0221** | **0.751** | **F1 Female (from Rejected vs Non-Rejected F0)** |
| **mml-miR-30a-3p** | **1.46** | **0.0252** | **0.751** | **F1 Female (from Rejected vs Non-Rejected F0)** |
| **mml-miR-133c-3p** | **-4.6** | **0.0258** | **0.751** | **F1 Female (from Rejected vs Non-Rejected F0)** |
| **mml-miR-20b-5p** | **2.19** | **0.0265** | **0.751** | **F1 Female (from Rejected vs Non-Rejected F0)** |
| **mml-miR-130b-3p** | **1.38** | **0.027** | **0.751** | **F1 Female (from Rejected vs Non-Rejected F0)** |
| **mml-miR-126** | **-1.58** | **0.0293** | **0.751** | **F1 Female (from Rejected vs Non-Rejected F0)** |
| **mml-miR-34a-5p** | **-1.93** | **0.0318** | **0.751** | **F1 Female (from Rejected vs Non-Rejected F0)** |
| **mml-miR-132-3p** | **1.5** | **0.0333** | **0.751** | **F1 Female (from Rejected vs Non-Rejected F0)** |
| **mml-let-7i-5p** | **-1.29** | **0.0389** | **0.812** | **F1 Female (from Rejected vs Non-Rejected F0)** |
| **mml-miR-320a** | **1.33** | **0.049** | **0.885** | **F1 Female (from Rejected vs Non-Rejected F0)** |

***Significantly upregulated, passing FDR correction**

**Supplementary File 8. Behaviours percentage of owl monkey parents with or without parental rejection**

| **Behaviour** | | **Parents with parental rejection (PR)**  N = 11^1^ | **Parents without parental rejection**  **(NPD)**  N = 27^1^ | **p-value**^2^ | **Average**  **Treatment Effect (PD-NPD)**^3^ |
| --- | --- | --- | --- | --- | --- |
| **Temerity** | Fear | 98 (0) | 95 (2) | 0.03 | **5 (2)** |
|  | Hidden in the nest | 4 (4) | 7 (6) | 0.09 | **-5 (3)** |
|  | Hidden in the hanger | 8 (8) | 84 (21) | <0.01 | **-44 (36)** |
|  | Hidden behind | 5 (6) | 9 (9) | 0.08 | **-11 (13)** |
|  | Frozen | 4 (3) | 8 (3) | 0.4 | **-8 (8)** |
| **Defecating** | Defecate | 6 (1) | 6 (4) | 0.4 | **7 (6)** |
| **Urinating** | Urinate | 8 (7) | 8 (7) | 0.7 | 0 (3) |
| **Bonding** | Cuddle | 96 (0) | 94 (1) | 0.01 | 2 (2) |
|  | Parent reaches offspring | 42 (28) | 68 (38) | 0.15 | 1 (2) |
|  | Carrying infants | 1 (2) | 7 (3) | 0.04 | **-7 (7)** |
| **Locomotion** | Locomotion jumping | 3 (0) | 7 (3) | <0.01 | **-5 (4)** |
|  | Locomotion walking* | 0/11 | 0/27 | - | - |
| **Stereotyping** | Head twirling | 6 (1) | 3 (3) | 0.15 | 1 (3) |
|  | Somersaults* | 0/11 | 0/27 | - | - |
|  | Pacing* | 1/11 | 0/27 | - | - |
| **Reactivity** | Furtive look | 94 (4) | 94 (5) | 0.2 | **-3 (3)** |
| **Audacity** | Approximation | 70 (41) | 6 (7) | 0.01 | **20 (14)** |
| **Curiosity** | Alert curiosity | 96 (0) | 96 (0) | 0.5 | -1 (3) |
| **Social Agonist** | Bite | 9 (5) | 33 (49) | 0.14 | **7 (9)** |
|  | Lance* | 1/11 | 0/27 | - | - |
|  | Attacks himself* | 0/11 | 0/27 | - | - |
|  | Hit others | 7 (1) | 3 (2) | 0.01 | **10 (9)** |

1 Median % (median absolute deviation %) adjusted by family/cage and sex effects using a logistic regression with bayesian estimation and uninformative priors

2 Wilcoxon rank sum exact test of adjusted values

3 Median treatment difference % (median absolute deviation %) estimated using the logistic regression adjusted by family/cage and sex effects and G-formula assigning all individuals to each treatment and estimating the difference between the 2 posterior distributions.

* Not enough individuals showing the behaviour to estimate any difference

**Supplementary file 9. Behaviours percentage of owl monkey mothers and fathers with or without parental deprivation**

| **Behaviour** | | **Mother with parental deprivation**  **(MD)***  N = 6^1^ | **Mother without parental deprivation (MND)***  N = 12^1^ | **p-value**^2^ | **Average**  **Treatment Effect (MD-MND)**^3^ | **Father with parental rejection**  **(FD)***  N = 5^1^ | **Father without parental rejection (FND)***  N = 15^1^ | **p-value**^2^ | **Average**  **Treatment Effect (FD-FND)**^3^ |
| --- | --- | --- | --- | --- | --- | --- | --- | --- | --- |
| **Temerity** | Fear | 98 (0) | 96 (1) | 0.01 | **4 (3)** | 95 (4) | 95 (1) | 0.7 | **5 (3)** |
|  | Hidden in the nest | 15 (20) | 5 (3) | 0.3 | **-5 (4)** | 4 (3) | 53 (68) | 0.14 | **-5 (3)** |
|  | Hidden in the hanger | 13 (17) | 82 (24) | 0.2 | **-44 (43)** | 3 (4) | 84 (23) | 0.01 | **-39 (23)** |
|  | Hidden behind | 39 (25) | 9 (2) | 0.9 | **-10 (8)** | 1 (0) | 38 (47) | <0.01 | **-11 (14)** |
|  | Frozen | 4 (0) | 10 (1) | 0.01 | **-10 (3)** | 6 (8) | 3 (1) | 0.9 | **-5 (4)** |
| **Defecating** | Defecate | 8 (2) | 6 (3) | 0.08 | **8 (6)** | 6 (0) | 4 (2) | 0.9 | **7 (6)** |
| **Urinating** | Urinate | 36 (42) | 8 (0) | 0.2 | -1 (2) | 3 (0) | 2 (0) | 0.7 | 1 (2) |
| **Bonding** | Cuddle | 96 (0) | 95 (1) | <0.01 | 2 (2) | 94 (4) | 94 (1) | 0.9 | 2 (2) |
|  | Parent reaches offspring | 41 (2) | 70 (36) | 0.8 | 1 (3) | 60 (54) | 68 (42) | 0.4 | 0 (2) |
|  | Carrying infants | 1 (1) | 2 (0) | 0.2 | **-4 (3)** | 4 (5) | 7 (2) | 0.03 | **-8 (3)** |
| **Locomotion** | Locomotion jumping | 3 (0) | 8 (3) | <0.01 | **-5 (4)** | 3 (0) | 7 (3) | <0.01 | **-5 (4)** |
|  | Locomotion walking* | 0/6 | 0/12 | - | - | 0/5 | 0/15 | - | - |
| **Stereotyping** | Head twirling | 6 (0) | 6 (1) | 0.9 | 1 (4) | 3 (0) | 3 (0) | 0.07 | 2 (3) |
|  | Somersaults* | 0/6 | 0/12 | - | - | 0/5 | 0/15 | - | - |
|  | Pacing* | 0/6 | 0/12 | - | - | 1/5 | 0/15 | - | - |
| **Reactivity** | Furtive look | 97 (0) | 98 (0) | <0.01 | -1 (2) | 94 (0) | 94 (0) | 0.9 | -3 (2) |
| **Audacity** | Approximation | 20 (19) | 1 (1) | 0.07 | **19 (18)** | 87 (14) | 6 (6) | 0.05 | **28 (13)** |
| **Curiosity** | Alert curiosity | 96 (0) | 96 (0) | 0.8 | 0 (3) | 96 (0) | 96 (0) | 0.5 | -1 (3) |
| **Social Agonist** | Bite | 10 (0) | 10 (0) | 0.3 | **13 (12)** | 6 (0) | 33 (48) | 0.6 | **4 (3)** |
|  | Lance* | 0/6 | 0/12 | - | - | 1/5 | 0/15 | - | - |
|  | Attacks himself* | 0/6 | 0/12 | - | - | 0/5 | 0/15 | - | - |
|  | Hit others | 7 (1) | 7 (1) | 0.07 | **11 (9)** | 5 (0) | 3 (2) | 0.07 | **8 (5)** |

1 Median % (median absolute deviation %) adjusted by family/cage and sex effects using a logistic regression with bayesian estimation and uninformative priors

2 Wilcoxon rank sum exact test of adjusted values

3 Median treatment difference % (median absolute deviation %) estimated using the logistic regression adjusted by family/cage and sex effects and G-formula assigning all individuals to each treatment and estimating the difference between the 2 posterior distributions.

* Not enough individuals showing the behaviour to estimate any difference

**Supplementary file 10. Behaviours percentage of owl monkey offspring from parents with or without parental deprivation**

| **Behaviour** | | **Female offspring from parents with parental deprivation**  **(FOD)***  N = 9^1^ | **Female offspring from parents without parental deprivation (FOND)***  N = 8^1^ | **p-value**^2^ | **Average**  **Treatment Effect (FOD-FOND)**^3^ | **Male offspring from parents with parental deprivation**  **(MOD)***  N = 5^1^ | **Male offspring from parents without parental deprivation (MOND)***  N = 5^1^ | **p-value**^2^ | **Average**  **Treatment Effect (MOD-MOND)**^3^ |
| --- | --- | --- | --- | --- | --- | --- | --- | --- | --- |
| **Temerity** | Fear | 92 (3) | 90 (5) | 0.2 | **18 (12)** | 94 (0) | 92 (1) | 0.4 | **10 (10)** |
|  | Hidden in the nest | 76 (35) | 7 (5) | 0.3 | **12 (11)** | 6 (1) | 5 (1) | 0.1 | **6 (5)** |
|  | Hidden in the hanger | 6 (3) | 95 (3) | 0.1 | **-15 (6)** | 6 (1) | 94 (1) | 0.3 | **-15 (3)** |
|  | Hidden behind | 59 (24) | 58 (59) | 0.7 | **15 (13)** | 5 (1) | 4 (6) | 0.4 | **4 (3)** |
|  | Frozen | 9 (2) | 9 (3) | 0.4 | **-11 (10)** | 7 (1) | 6 (2) | 0.3 | -3 (4) |
| **Defecating** | Defecate | 6 (5) | 6 (1) | 0.7 | **-6 (7)** | 5 (0) | 6 (1) | 0.03 | -2 (6) |
| **Urinating** | Urinate | 6 (1) | 5 (1) | 0.8 | **-5 (6)** | 7 (1) | 7 (2) | 0.7 | **-7 (6)** |
| **Bonding** | Cuddle | 94 (5) | 94 (3) | 0.3 | -1 (6) | 94 (0) | 94 (2) | >0.9 | -1 (5) |
|  | Carrying siblings | 7 (1) | 7 (1) | 0.2 | **-6 (5)** | 5 (0) | 6 (2) | 0.5 | -1 (4) |
| **Locomotion** | Locomotion jumping | 5 (1) | 6 (1) | <0.01 | **-14 (6)** | 5 (0) | 7 (3) | 0.2 | **-14 (1)** |
|  | Locomotion walking* | 0/9 | 0/8 | - | - | 0/5 | 0/5 | - | - |
| **Stereotyping** | Head twirling* | 0/9 | 0/8 | - | - | 0/5 | 0/5 | - | - |
|  | Somersaults* | 0/9 | 0/8 | - | - | 0/5 | 0/5 | - | - |
|  | Pacing* | 0/9 | 0/8 | - | - | 0/5 | 0/5 | - | - |
| **Reactivity** | Furtive look | 95 (0) | 94 (0) | 0.06 | **7 (2)** | 94 (0) | 96 (0) | 0.8 | 3 (6) |
| **Audacity** | Approximation | 7 (5) | 6 (3) | 0.9 | 3 (6) | 7 (0) | 6 (7) | 0.1 | 1 (5) |
| **Curiosity** | Alert curiosity | 95 (1) | 94 (1) | 0.2 | 4 (3) | 94 (0) | 95 (1) | 0.7 | 1 (6) |
| **Social Agonist** | Bite | 5 (1) | 17 (17) | 0.01 | **-11 (4)** | 7 (0) | 7 (2) | 0.3 | **-12 (12)** |
|  | Lance* | 0/9 | 0/8 | - | - | 0/5 | 0/5 | - | - |
|  | Attacks himself* | 0/9 | 0/8 | - | - | 0/5 | 0/5 | - | - |
|  | Hit others* | 0/9 | 0/8 | - | - | 0/5 | 0/5 | - | - |

1 Median % (median absolute deviation %) adjusted by family/cage, age category, and sex effects using a logistic regression with bayesian estimation and uninformative priors

2 Wilcoxon rank sum exact test of adjusted values

3 Median treatment difference % (median absolute deviation %) estimated using the logistic regression adjusted by family/cage, age category, and sex effects, and G-formula assigning all individuals to each treatment and estimating the difference between the 2 posterior distributions.

* Not enough individuals showing the behaviour to estimate any difference

**Supplementary file 11. Behaviours percentage of owl monkey offspring from parents with or without parental rejection**

| **Behaviour** | | **Offspring from one parent with parental deprivation (OPD)**  N = 14^1^ | **Offspring from parents without parental deprivation**  **(ONPD)**  N = 13^1^ | **p-value**^2^ | **Average**  **Treatment Effect (OPD-ONPD)**^3^ |
| --- | --- | --- | --- | --- | --- |
| **Temerity** | Fear | 93 (3) | 91 (3) | 0.14 | **18 (13)** |
|  | Hidden in the nest | 21 (26) | 6 (3) | 0.13 | **12 (10)** |
|  | Hidden in the hanger | 6 (3) | 94 (3) | 0.05 | **-15 (5)** |
|  | Hidden behind | 51 (32) | 24 (34) | 0.8 | **9 (10)** |
|  | Frozen | 8 (3) | 8 (2) | 0.9 | **-7 (8)** |
| **Defecating** | Defecate | 5 (2) | 6 (1) | 0.14 | **-6 (8)** |
| **Urinating** | Urinate | 6 (1) | 6 (1) | 0.7 | **-5 (5)** |
| **Bonding** | Cuddle | 94 (3) | 94 (3) | 0.4 | -1 (6) |
|  | Carrying siblings | 5 (2) | 7 (2) | 0.2 | **-6 (6)** |
| **Locomotion** | Locomotion jumping | 5 (0) | 7 (1) | <0.01 | **-14 (4)** |
|  | Locomotion walking* | 0/14 | 0/13 | - | - |
| **Stereotyping** | Head twirling* | 0/14 | 0/13 | - | - |
|  | Somersaults* | 0/14 | 0/13 | - | - |
|  | Pacing* | 0/14 | 0/13 | - | - |
| **Reactivity** | Furtive look | 95 (1) | 94 (0) | 0.3 | **7 (3)** |
| **Audacity** | Approximation | 7 (3) | 6 (3) | 0.3 | 3 (6) |
| **Curiosity** | Alert curiosity | 95 (1) | 94 (1) | 0.7 | **4 (3)** |
| **Social Agonist** | Bite | 6 (1) | 7 (3) | 0.01 | **-12 (4)** |
|  | Lance* | 0/14 | 0/13 | - | - |
|  | Attacks himself* | 0/14 | 0/13 | - | - |
|  | Hit others* | 0/14 | 0/13 | - | - |

1 Median % (median absolute deviation %) adjusted by family/cage, age category, and sex effects using a logistic regression with bayesian estimation and uninformative priors

2 Wilcoxon rank sum exact test of adjusted values

3 Median treatment difference % (median absolute deviation %) estimated using the logistic regression adjusted by family/cage, age category, and sex effects, and G-formula assigning all individuals to each treatment and estimating the difference between the 2 posterior distributions.

* Not enough individuals showing the behaviour to estimate any difference
